# Unisexual reproduction in the global human fungal pathogen *Cryptococcus neoformans*

**DOI:** 10.1101/2025.06.02.657540

**Authors:** Sheng Sun, Zhuyun Bian, Ziyan Xu, Yeseul Choi, Joseph Heitman

## Abstract

The human fungal pathogen *Cryptococcus* species complex (encompassing *C. neoformans*, *C. deneoformans*, and the *C. gattii* species complexes) exhibit diversity in sexual reproduction, including α-**a** mating, pseudosexual reproduction, as well as unisexual reproduction initiated from a single isolate or between isolates of the same mating type. A central conundrum is that while most *Cryptococcus* natural populations exhibit significant α mating-type bias, genetic and genomic analyses show recombination occurs in nature. The discovery of unisexual reproduction in *C. deneoformans* provided insight; however, thus far unisexual reproduction has never been directly observed in the predominant global pathogenic species *C. neoformans*. Here, we provide evidence that mutating the *RIC8* gene, which encodes a conserved guanine nucleotide exchange factor (GEF) involved in both chaperoning and activating Gα proteins, enables unisexual reproduction in *C. neoformans*. Additionally, we show that genetic variation in the natural population promotes unisexual reproduction, and unisexual reproduction in *C. neoformans* involves canonical meiotic recombination. Finally, we found that deletion of both *GPA2* and *GPA3* in the *MAT*α background leads to self-filamentation without sporulation, suggesting that differential modulation of the Gα proteins, likely involving Ric8, could underlie the switch between different modes of sexual reproduction in *Cryptococcus*. Our study further highlights that the highly conserved Ric8 GEF can act as an important regulator of cellular development in response to environmental stimuli and could modulate sexual reproduction in nature. We hypothesize that unisexual reproduction occurs much more frequently in nature than currently appreciated, and possibly in other fungi and microbial eukaryotes as well.

**Significance statement:** Sexual reproduction is a central tenet of the eukaryotic life cycle, essential for the long-term survival of a species by promoting genetic diversity and facilitating natural selection. In eukaryotes such as unicellular basidiomycetous fungi, sexual reproduction leads to hyphal growth and spore production, enabling escape from harsh environments and long-distance dispersal. Here, we report that mutating the *RIC8* gene, encoding a conserved guanine nucleotide exchange factor (GEF) involved in activation of Gα proteins, enables unisexual reproduction in the predominant human fungal pathogen *Cryptococcus neoformans* that involves canonical meiosis. Our findings suggest that conditions might exist in nature that could induce, and even promote, unisexual reproduction in *Cryptococcus*, highlighting the complexity and importance of sexual reproduction in this prominent fungal pathogen.

## Introduction

Sexual reproduction is a fundamental process in eukaryotes. Through meiosis, sex can produce progeny populations with increased genetic diversity and phenotypic variation, therefore facilitating natural selection and playing a critical role in the long-term survival of a species. Sexual reproduction is of particular importance to unicellular eukaryotes such as fungi. While most fungal species typically reproduce mitotically in nature, they are susceptible to changing environments such as fluctuations in temperature, nutrients, antifungal natural products, as well as co-inhabitants in their natural niches. Sexual reproduction allows generation of recombinant progeny with higher fitness in the current environment, as well as production of sexual spores that are better equipped for dispersal and withstanding harsh environments. Additionally, for yeast species, sexual reproduction can be linked with hyphal growth, which can facilitate foraging for nutrients and escaping from unfavorable conditions. Fungi exhibit extraordinary diversity in the modes of sexual reproduction, including α-**a** mating, unisexual reproduction, parasexual reproduction, and pseudosexual reproduction (1–9).

One example of this reproductive plasticity can be found in the human fungal pathogen *Cryptococcus* species complex, which are basidiomycetous yeasts that cause pulmonary infections, as well as systemic infections leading to meningoencephalitis. Composed of *C. neoformans*, *C. deneoformans*, and six species within the *C. gattii* species complex, the pathogenic *Cryptococcus* species collectively cause over 220,000 infections globally each year, accounting for ∼181,000 deaths annually (10). There are two mating types in *Cryptococcus*, α and **a**, defined by a single mating type locus (*MAT*) in the genome, and sexual reproduction between α and **a** cells of *Cryptococcus* (α-**a** sexual reproduction) was first characterized in the laboratory ∼50 years ago (11–13). Studies have focused on modes of sexual reproduction because spores, which are readily aerosolized and of an optimal size for alveolar deposition following inhalation, are documented infectious propagules (14–16).

Among natural *Cryptococcus* isolates, there is a significant bias toward the α mating type (>99.9% in some cases), and populations with more balanced α and **a** mating types have only been found in a few restricted geographic areas (e.g. sub-Saharan Africa) (17). Nevertheless, population genetics and genomics studies indicate that most natural populations of *Cryptococcus* are recombining, suggesting sexual reproduction is ongoing, even in populations with few, or no, *MAT***a** isolates (18, 19). How a recombining population with highly unbalanced mating types is maintained has long puzzled researchers.

Some 20 years ago, Lin et al. (20) discovered a novel mode of sexual reproduction in *C. deneoformans*, termed unisexual reproduction. In this case, sexual reproduction can be initiated by either a single cell (through endoreplication) or two cells of the same mating type (through conjugation/mating) and then undergo sexual development similar to α-**a** sexual reproduction, with hyphal growth, basidium formation, meiosis, and sporulation (21–25). Subsequently, unisexual reproduction has been characterized in other fungal species such as *Candida albicans* (26). Unisexual reproduction could potentially explain the observed mating type imbalance observed in *Cryptococcus* natural populations. However, until now it has only been well characterized in *C. deneoformans*. Recently, we showed that in *C. deneoformans* Ric8 is involved in regulating cell morphological development, and mutation of the *RIC8* gene enables cells to undergo titanization and unisexual development in solo cultures under mating inducing conditions (27). Interestingly, we also discovered that the *C. deneoformans* strain XL280, which undergoes robust unisexual reproduction (20), harbors a nonsense mutation in the *RIC8* gene, indicating that Ric8 suppresses unisexual reproduction (27).

Ric8 (resistance to inhibitors of cholinesterase 8) and its homologs are conserved in a wide range of eukaryotes, from unicellular fungi to invertebrates and mammals. Functioning as both a non-receptor guanine nucleotide exchange factor (GEF) and a molecular chaperone, Ric8 catalyzes the exchange of GDP for GTP on Gα subunits and promotes their proper folding and membrane association (28–32). In contrast to G protein-coupled receptors (GPCRs), Ric8 lacks a transmembrane domain and operates entirely within the cytoplasm to modulate Gα activity (33, 34), similar to a group of other nonreceptor proteins that contain the “Gα-binding and -activating (GBA) motif” (35). In fungi, Ric8 regulates G-protein signaling required for the nematode-trapping lifecycle of *Arthrobotrys oligospora* (36). In animals, Ric8 functions as an intracellular protein that plays a pivotal role in the regulation of heterotrimeric G protein signaling, and is involved in a variety of key cellular processes such as cell polarity establishment, cytokinesis, and nervous system development, underscoring its essential role in embryogenesis and neurodevelopment (37). For example, in *Caenorhabditis elegans* Ric8 acts as a regulator of asymmetric cell division and centrosome dynamics (38). In mammals, Ric8 is expressed in two functionally distinct isoforms, Ric8A and Ric8B, that exhibit selective binding and activation profiles toward specific Gα subsets, and regulate distinct G protein signaling pathways, with Ric8A being associated with embryonic development and central nervous system formation through Gα_i_ and Gα_q_ signaling, and Ric8B involved in olfactory signaling and neural development via activation of Gα_s/olf_ (32–34, 39). Despite the partial overlap in their target Gα subunits, Ric8A and Ric8B serve non-redundant and essential functions in vivo, with complete deletion of either gene resulting in embryonic lethality in mice. Additionally, Ric8A and Ric8B deletion result in severe depletion of specific Gα proteins, suggesting that both isoforms are required for proper Gα protein maturation (34).

In *Cryptococcus*, studies have provided evidence that Ric8 interacts physically and functionally with two of the three Gα proteins, Gpa1 and Gpa2, but not Gpa3, and plays an important role in regulation of virulence related traits such as melanin and capsule production, as well as response to mating pheromones (40), and is required for titanization in *C. neoformans* (41). It is possible that Ric8, functioning through binding and activating the Gα proteins, could influence conserved GPCR pathways that transmit environmental stimuli, thus modulating the downstream signaling pathways involved in morphological development in *Cryptococcus*. This is consistent with previous study demonstrating that Ric8 is involved in modulating virulence potential, high temperature tolerance, and melanin production (42), as well as our findings that Ric8 represses morphological development (selfing and titanization) in *C. deneoformans* (27). We hypothesized that Ric8 plays a similar role in repressing morphological development, including unisexual reproduction, in *C. neoformans*. Here, we show that mutating the *RIC8* gene induces the *C. neoformans* lab reference type strain H99 to undergo unisexual reproduction. Additionally, we generated strains with enhanced unisexual fertility by crossing H99 *ric8* mutants with natural isolates and isolating F1 progeny with enhanced unisexual fertility, revealing the presence of variation promoting unisexual reproduction in nature. With these robust unisexually fertile strains, we elucidated the signaling pathways required for unisexual reproduction in *C. neoformans* and show that sporulation during *C. neoformans* selfing involves canonical meiosis. Our findings further highlight the diverse approaches that pathogenic *Cryptococcus* species utilize to accomplish sexual reproduction, with implications for their ecology, population genetics, epidemiology, and evolution.

## Results

### Deletion of *RIC8* enables unisexual reproduction in *C. neoformans* lab strains

In a recent study, we discovered that Ric8, a guanine nucleotide exchange factor (GEF), acts as a suppressor of morphological development (e.g. titan cells and unisexual reproduction) in *C. deneoformans* under conditions inducing unisexual reproduction (27). Interestingly, the well characterized *C. deneoformans* isolate XL280, which undergoes highly robust unisexual reproduction, has a nonsense mutation in the *RIC8* gene, rendering it non-functional. Taking this lead, we hypothesized that by deleting the *RIC8* gene, unisexual reproduction might be induced in the predominant global pathogenic species *C. neoformans*. To test this, the *RIC8* gene was deleted in the *C. neoformans* lab strain KN99α to generate KN99α *ric8*11::*NAT* mutant strains (SSI867 and SSI868, Supplemental Table S1). When grown under mating inducing conditions, structures resembling sexual development, including hyphae, basidia, and basidiospores were observed (Figure 1). Basidiospores were dissected from five individual basidia produced by strain SSI867 and analyses revealed: 1) spores from all of the basidia had high germination rates, 2) all of the spores harbored the NAT-resistant *RIC8* deletion, and 3) all of the basidiospores were *MAT*α (Table 1). These findings support the conclusion that these basidiospores were all produced by the *MAT*α *ric8* NAT-resistant strain (i.e. SSI867) through unisexual reproduction.

**Figure 1.**
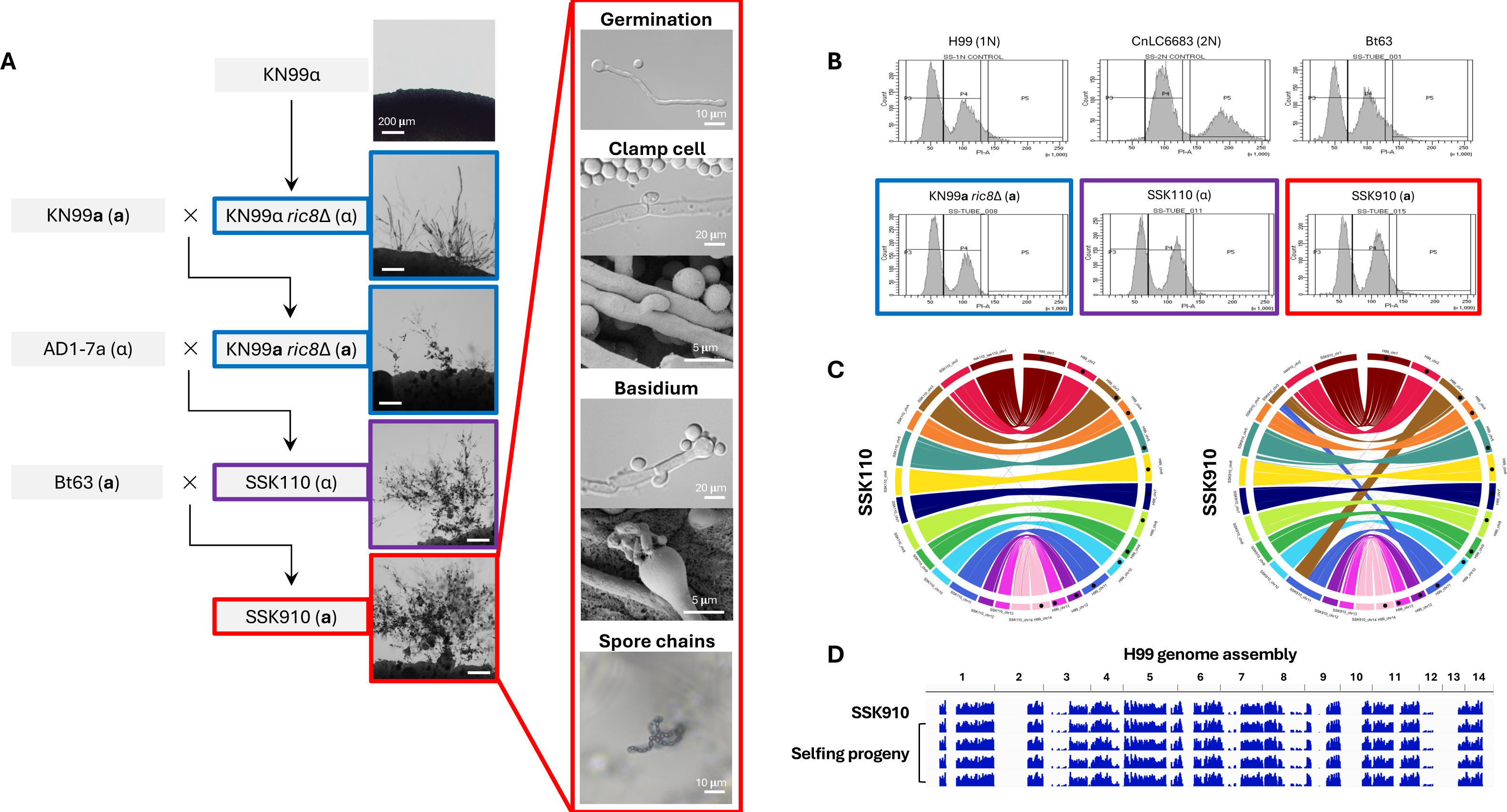
Haploid *C. neoformans* strains that undergo robust unisexual reproduction. (A) Genealogy of strains involved in constructing *C. neoformans* isolates that undergo robust unisexual reproduction, with light microscopy images next to the strains illustrating sexual structures on MS medium. Highlights with blue, purple, and red boundaries indicate low, medium, and high unisexual development, respectively. For strain SSK910 that undergoes the most robust unisexual reproduction, additional images for germination tube, unfused clamp cell, basidium, and basidiospore chains are provided. (B) Flow cytometry analysis reveals unisexually fertile strains are haploid. (C) Circos plots of strains SSK110 (left) and SSK910 (right) reveal their genomes are colinear with the *C. neoformans* reference strain H99. For each plot, the right half shows chromosomes 1 to 14 (clockwise) of strain H99, and the left chromosomes 1 to 14 (counterclockwise) of strains SSK110 or SSK910. The discrepancies between strains SSK910 and H99 involving chromosomes 3 and 11 correspond to the reciprocal translocation between the two chromosomes in H99 as reported previously (69). (D) WGS and variant analysis of strain SSK910 and basidiospores dissected from SSK910 are depicted. Sequencing reads were mapped, and variants called with strain H99 as reference; blue bars indicating sequences that differ from H99. Both SSK910 and its progeny have identical genomes that are recombinant between H99 and non-H99 sequences (inherited from the natural isolates AD1-7a and Bt63 during construction of strain SSK910).

**Table 1.**
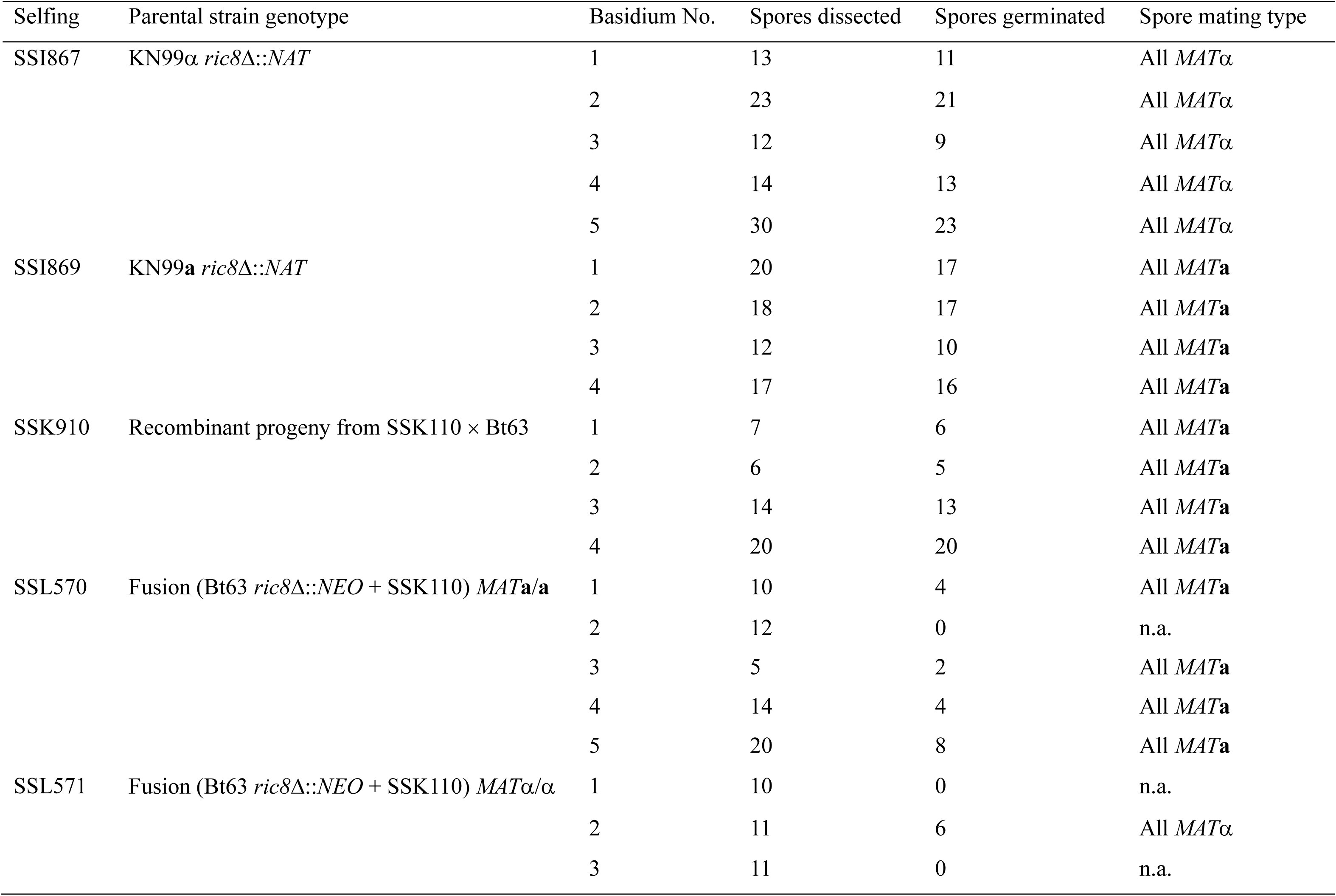
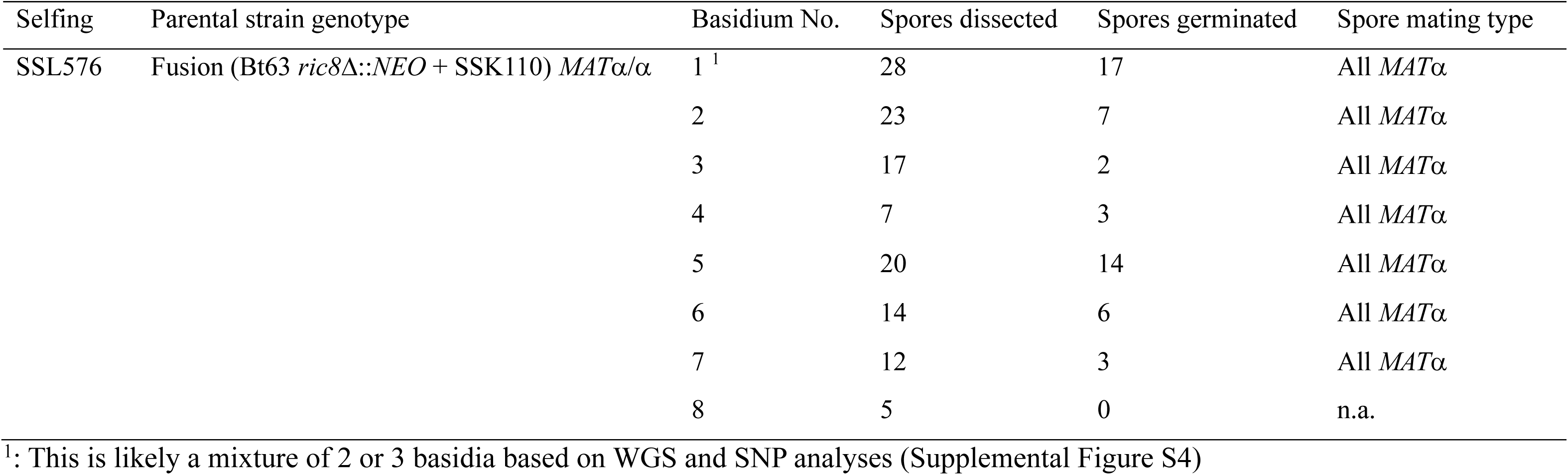
Unisexual reproduction and progeny sets generated and analyzed in this study. .

Next the KN99α *ric8* mutants (SSI867 and SSI868) were crossed with strain KN99**a** that is congenic with KN99α, and *MAT***a** *ric8*11::*NAT* progeny (SSI869 and SSI870, Supplemental Table S1) were recovered. Similar to the KN99α *ric8* mutant strains SSI867 and SSI868, solo cultures of the *MAT***a** *ric8*11::*NAT* progeny SSI869 and SSI870 underwent sexual development under mating inducing conditions (Figure 1). Analyses of four individual basidia produced by SSI869 revealed high germination rates in the basidiospores and showed that they were all *MAT***a** *ric8*11::*NAT*, again consistent with spore production via unisexual reproduction of the *MAT***a** *ric8* strain SSI869.

Taken together, these data indicate that deletion of the *RIC8* gene in the *C. neoformans* lab strains KN99α and KN99**a** enables unisexual reproduction, consistent with our previous findings in the sister species *C. deneoformans* (27).

### Naturally occurring variation promotes unisexual reproduction in *C. neoformans*

While the *C. neoformans* lab strains with a *ric8*11::*NAT* deletion are able to undergo unisexual reproduction, it occurs at relatively low frequencies and often requires prolonged incubation (:= 4 weeks) under mating inducing conditions. We hypothesized that polymorphisms existing in natural isolates might promote unisexual reproduction. To test this, strain SSI867 (KN99α *ric8*11::*NAT*) was crossed with two *MAT***a** natural isolates (Bt63 and Bt65); strain SSI869 (KN99**a** *ric8*11::*NAT*) was crossed with two *MAT*α natural isolates (AD-17a and T4) (Table 2). A total of 56 random spores were dissected from each of these four crosses, with germination rates ranging from 18% (SSI867 × Bt65) to 34% (SSI869 × AD1-7a) (Table 2). Indeed, with the exception of the cross between SSI869 and T4, progeny were recovered with comparable or enhanced unisexual fertility compared to the unisexual parent (i.e. SSI867 or SSI869) in the other three crosses, at a percentage between 10% (3 out of 30 progeny, SSI867 × Bt63) and 30% (3 out of 10 progeny, SSI867 × Bt65) (Table 2). Interestingly, all of the unisexually fertile progeny inherited the *ric8*11::*NAT* allele from their selfing parent, further supporting that the ability to undergo unisexual reproduction requires the *ric8* mutation (Table 2).

**Table 2.**
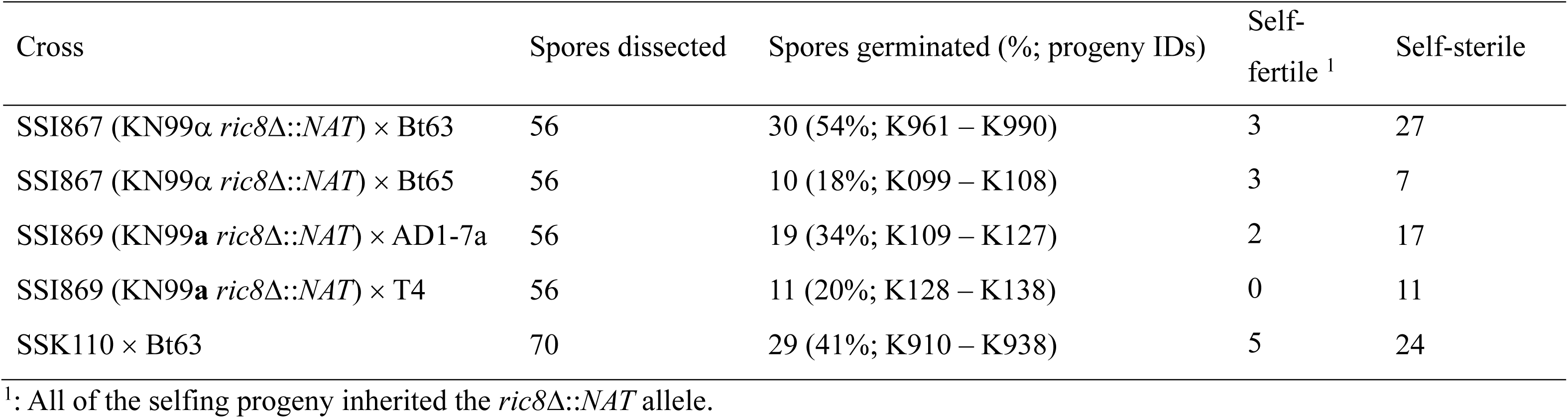
α-a sexual crosses and progeny sets generated and analyzed in this study. .

Of the unisexually fertile progeny that were recovered, one (SSK110) from the SSI869 × AD1-7a cross showed considerably enhanced selfing ability (Figure 1A and Table 2). SSK110 is a haploid *MAT*α strain (Figure 1B), with a genome that is colinear with the *C. neoformans* reference strain H99 (Figure 1C). Strain SSK110 was further crossed with natural isolate Bt63 (VNBI, *MAT***a**) and 70 random spores were dissected. Of the 29 progeny that germinated, five were able to self, with one progeny (SSK910) undergoing very robust unisexual reproduction under mating inducing conditions, producing extensive hyphae with unfused clamp cells and abundant basidia and basidiospore chains (Figure 1A). Strain SSK910 is a haploid *MAT***a** strain (Figure 1B), and its genome is nearly colinear with the reference strain H99, except for the reciprocal translocation involving chromosomes 3 and 11 that has been previously reported in the H99 genome (Figure 1C).

Self-fertile progeny (SSK910) from cross between SSK110 and Bt63 was further induced to undergo unisexual reproduction and random basidiospores were dissected. Whole genome sequencing and SNP analyses with H99 as reference showed that strain SSK910 and its F1 progeny have identical genomes (Figure 1D), which are recombinants of alleles from KN99 (regions with no SNPs) and natural isolates (i.e. AD1-7a and Bt63, regions in blue indicating variants against H99) across the genome (Figure 1D).

In summary, through intercrossing with natural isolates, we successfully generated *C. neoformans* strains that undergo robust unisexual reproduction, confirming the presence of natural variants that promote selfing.

### Unisexual reproduction in *C. neoformans* requires a functional MAPK pathway

With the *C. neoformans* strain SSK910 that undergoes robust unisexual reproduction, we sought to investigate the signaling pathways required for this process. We constructed deletion strains for 14 genes that have been previously shown to be involved in mating in *C. neoformans* in the SSK910 background, including those involved in G-protein signaling, mating pheromone sensing, the MAP kinase pathway, sexual development, and meiosis (Supplemental Table S2). We then studied whether these deletion strains were impaired for unisexual reproduction, including hyphal growth, basidium formation, and basidiospore production.

Among the 14 genes tested, seven were not required for unisexual reproduction, and they are involved in G-protein signaling (*GPA1*, *GPA2*, *GPA3*), PKA signaling (*PKA1*), mating pheromone or nutrient sensing (*STE3* and *GPR4*), and the α-**a** sexual reproduction specific transcription factor (*SXI2***a**) (Figure 2 and Supplemental Table S2) (40, 43–45). It should be noted that the deletions of *GPA1* and *PKA1* resulted in noticeably delayed unisexual reproduction, although in these cases, hyphae, basidia, and basidiospore chains were eventually produced within the selfing patches. Five genes were essential for unisexual reproduction, as deletion of these genes led to complete abolition of selfing, and the deletion strains grew only as yeast cells under mating inducing conditions (Figure 2). Three of these five genes (*CPK1*, *STE7*, and *STE11***a**) are components of the pheromone responsive MAP kinase signaling pathway (46), while the other two (*MAT2* and *ZNF2*) are known master regulator transcription factors for α-**a** sexual reproduction in *C. neoformans* and unisexual reproduction in *C. deneoformans* (47). Additionally, while strains deleted for *SPO11* or *DMC1*, two genes known to be required for meiosis (48, 49), were able to undergo the initial stages of unisexual development, producing hyphae and basidia, the production of basidiospore chains was severely impaired, consistent with compromised meiosis in these mutant strains (Figure 2 and Supplemental Figure S6).

**Figure 2.**
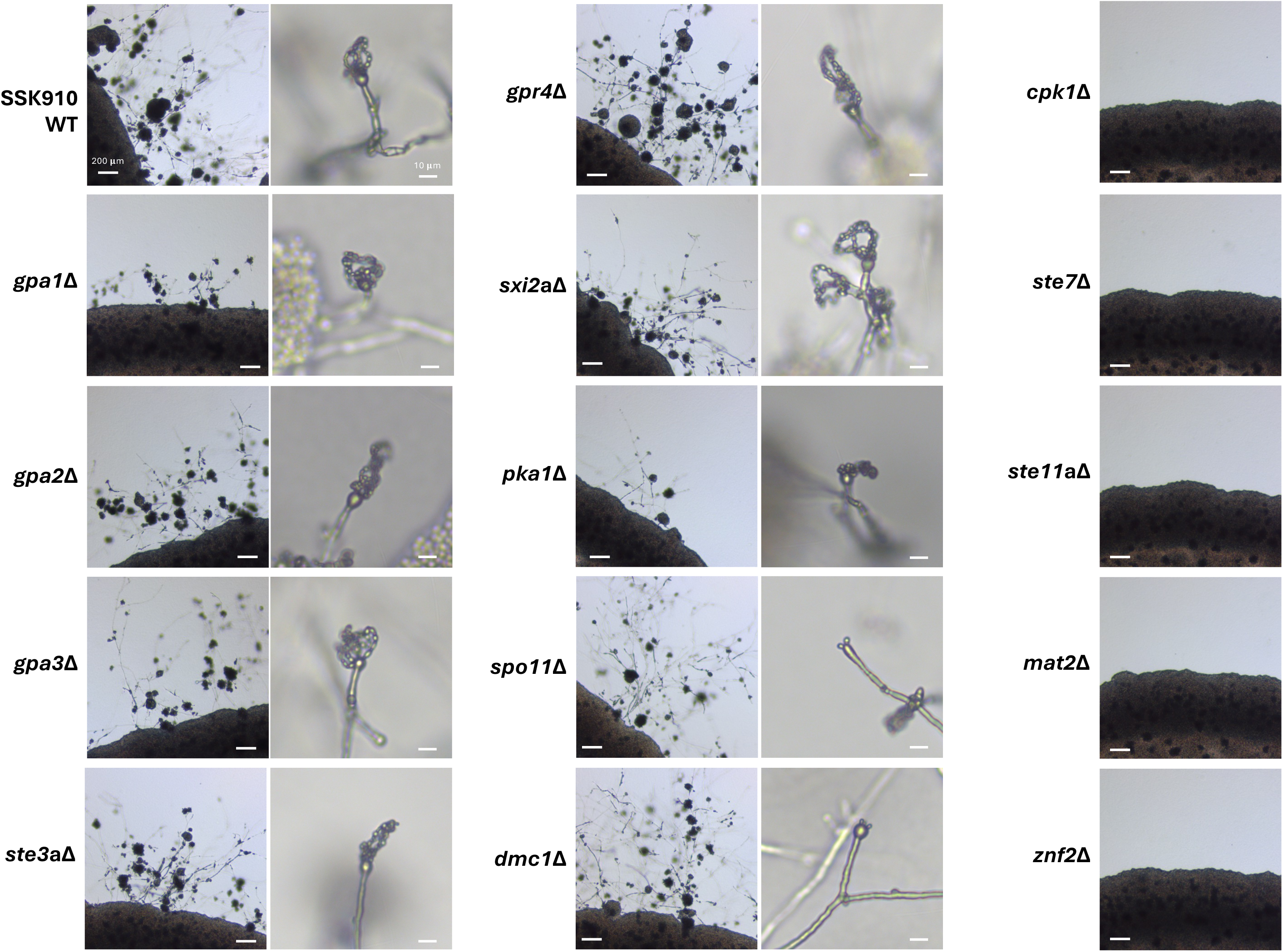
Unisexual reproduction in *C. neoformans* is dependent upon the MAPK pathway. Light microscopy images of strains with deletions of various genes in the SSK910 background when induced for unisexual reproduction on MS medium. Images shown here are from one of at least two independent deletion strains of each gene. Images on the right showed the basidia with or without basidiospore chains for the deletion strains that were still able to undergo unisexual reproduction.

Taken together, these results show that while the genes involved in G-protein signaling and mating pheromone sensing are dispensable or redundant, successful unisexual reproduction in *C. neoformans* requires a functional MAP kinase pathway, the master regulators of sexual reproduction, and genes essential for meiosis.

### Meiotic recombination occurs during unisexual reproduction in *C. neoformans*

Our results from gene deletion analyses show that unisexual reproduction in *C. neoformans* involves genes required for meiosis. We next investigated whether unisexual reproduction in *C. neoformans* involves canonical meiotic recombination and if so, how this compares to α-**a** sexual reproduction. To accomplish this, one approach could be to analyze progeny from crosses between two genetically divergent strains of the same mating type. However, this approach is not feasible for a couple of reasons. First, sexual reproduction between two *C. neoformans* wild-type isolates of the same mating type has not yet been observed and may occur at a low frequency due to bottlenecks such as cell-cell fusion. Second, it is possible that sexual reproduction could occur between two divergent *C. neoformans ric8*11::*NAT* isolates that each could undergo unisexual reproduction. However, in this case it would not be possible to differentiate morphologically between spores produced by selfing of each parent strain and those generated by crossing between the two parental isolates, thus complicating the analyses of meiotic products, because those from selfing lack genetic polymorphisms and thus are not suitable for analyzing meiotic recombination.

To overcome this, we first generated diploid *MAT***a**/**a** and *MAT*α/α strains that are homozygous for the *ric8*11/*ric8*11 deletion (enabling selfing) and also contain heterozygous polymorphisms throughout the genome (allowing analyses of allele segregation). Specifically, we generated a *ric8*11::*NEO* deletion strain in the natural isolate Bt63 background (SSL436). Next, we fused strain SSL436 (*MAT***a** *ric8*11::*NEO*) with SSK110 (*MAT*α *ric8*11::*NAT*), a self-fertile recombinant progeny from the cross between lab strain KN99**a** *ric8*11::*NAT* and natural isolate AD1-7a and recovered fusion products that were resistant to both *NAT* and *NEO*. We then screened for fusion isolates that also exhibited reduced self-fertile sexual reproduction (<1%) when compared to *C. neoformans MAT***a** × *MAT*α sexual mating controls under mating inducing conditions, indicative of loss of heterozygosity at the *MAT* locus resulting in *MAT***a**/**a** or *MAT*α/α strains undergoing unisexual reproduction. By this approach, we successfully isolated three diploid fusion products between strains SSK110 and SSL436 that are *MAT*-homozygous and genome-heterozygous: SSL570 (*MAT***a**/**a**), SSL571 (*MAT*α/α), and SSL576 (*MAT*α/α) (Figure 3A and Supplemental Table S1). Flow cytometry analyses confirmed that these fusion products are diploid (Figure 3B) and whole genome sequencing and variant analyses further showed that these strains contain heterozygosity throughout their genomes (Supplemental Figure S1, variants against H99 genome indicated in blue and variants against the Bt63 genome highlighted in red).

**Figure 3.**
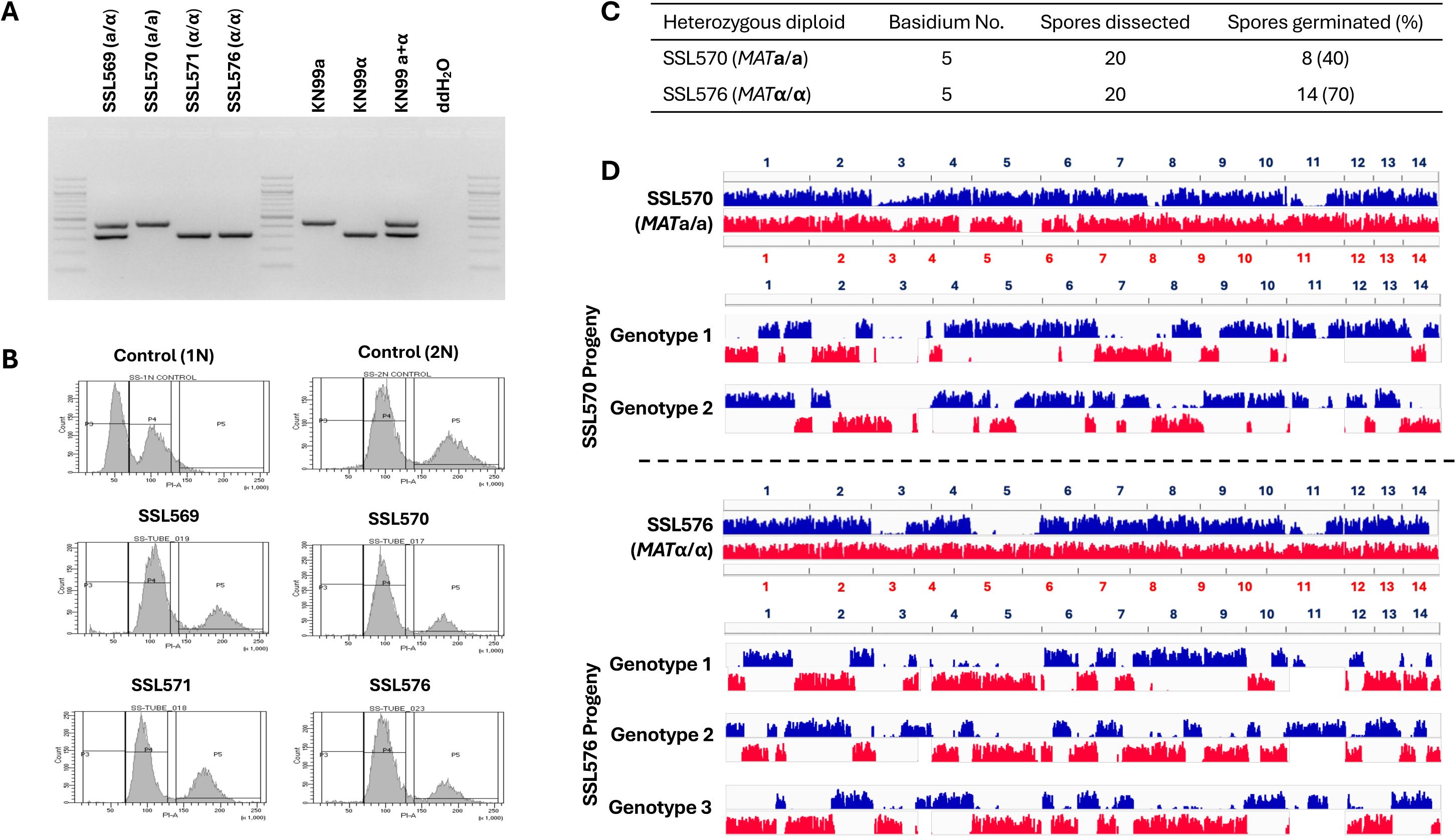
Unisexual reproduction in *C. neoformans* involves meiotic recombination. (A) Genotyping of the *MAT* locus showed that the three derived diploid strains, SSL570, SSL571, and SSL576, all underwent LOH at the mating type locus. (B) Flow cytometry analysis showed that the three derived diploid strains all have a diploid ploidy profile. (C) Summary of two basidia dissected from strains SSL570 and SSL576, respectively. (D) Variant mapping of the parental strains and their respective basidiospores. For the parental strains SSL570 and SSL576, the variants against H99 (blue) and Bt63 (red) were mapped along the H99 and Bt63 genomes, respectively. For the progeny, the variants against H99 and Bt63 were all mapped with the H99 genome as reference.

We then induced unisexual reproduction in diploid strains SSL570, SSL571, and SSL576 under mating inducing conditions, and dissected spores from individual basidia (Table 1) from each strain. Compared to the basidiospores dissected from unisexual reproduction of haploid *ric8*11 strains (e.g. SSI876 and SSK910, Table 1), the germination rates of spores from selfing of the diploid *MAT*-homozygous strains were considerably lower, which is reflective of both the genetic diversity and one chromosomal translocation between the two nuclear genomes in the diploids. This is consistent with the reduced spore germination rates that have been observed previously in crosses between *C. neoformans* lab strains and natural isolates (42, 50, 51). It should be noted that all of the progeny share the same mating type as their parental diploid strains, further confirming that these progeny are products of unisexual reproduction (Table 1).

Whole-genome-sequencing and variant analyses of the progeny dissected from these *MAT*-homozygous diploids shows clear evidence of allele segregation consistent with meiotic recombination (Figure 3D, Supplemental Figures S2 – S4). For example, of the 20 spores dissected from basidium No. 5 of strain SSL570 (*MAT***a**/**a**), eight germinated, which is a 40% germination rate (Figure 3C). Because in *C. neoformans* only one round of meiosis occurs within each basidium, giving rise to four recombinant meiotic products that undergo repeated rounds of mitosis to produce basidiospores (52), we would expect two distinct nuclear genotypes among these eight germinated spores given the ∼50% germination rate, assuming the other two genotypes failed to germinate due to the factors such as the known chromosomal translocations, as well as the segregation of other genetic polymorphisms between the two nuclear genomes. Indeed, we observed two types of progeny among these eight spores, each having recombinant genomes with alleles from the two parental nuclei alternating along each chromosome, consistent with products of meiotic crossovers (Figure 3D, top panel). Similarly, 14 of the 20 spores dissected from strain SSL576 basidium No. 5 germinated, and the 70% germination rate suggested that three of the four meiotic products gave rise to viable basidiospores. Indeed, we observed three different genotypes among these progeny (Figure 3D, bottom panel). Similar to those from SSL570 basidium No. 5, these progeny also possessed recombinant chromosomes that were consistent with meiotic crossovers. Additionally, no allele from either of the two parental genomes was observed in more than two progeny genotypes, providing further support that these progeny are products of meiosis.

Collectively, these analyses provided robust evidence that meiotic recombination occurs during unisexual reproduction in *C. neoformans*.

### Deletion of the Gpa2 and Gpa3 proteins leads to self-filamentation

Of the three Gα proteins in *C. neoformans*, Ric8 has been previously shown to interact with Gpa1 and Gpa2, but not Gpa3, based on yeast two hybrid assays and co-immunoprecipitation analysis (40). Interestingly, when we predicted protein-protein interactions between Ric8 and the three Gα proteins using AlphaFold multimer, our results suggest that Ric8 might form complexes with each Gα protein with similar interaction profiles of comparable confidence (Figure 4A).

**Figure 4.**
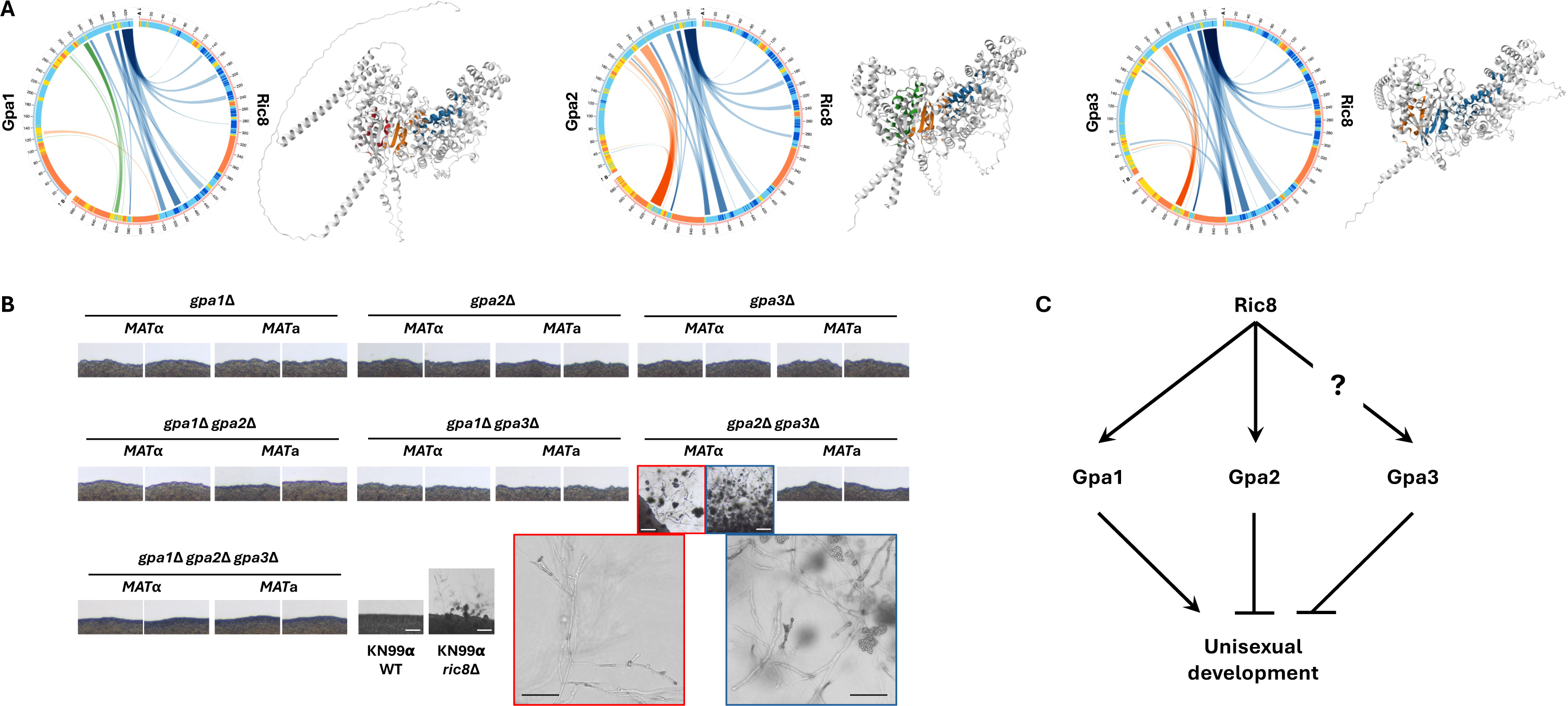
Genetic interactions between Ric8 and the three Gα proteins in *C. neoformans*. (A) AlphaFold predictions of the 3D structures and protein-protein interaction profiles between Ric8 and the three Gα proteins (Gpa1, Gpa2, and Gpa3). For each Ric8-Gα protein pair, the circus plot on the left illustrates the residues predicted to be interacting between the two proteins, and the image on the right shows the predicted 3D structure of the protein complex, colored for the interfaces. (B) Selfing assay of solo cultures of single, double, and triple deletion strains of the three Gα proteins. While most of the deletion strains assayed were self-sterile and grew as yeasts under mating inducing conditions, the two *MAT*α *gpa2*11 *gpa3*11 double deletion strains were self-filamentous and produce robust hyphae in solo cultures. However, no basidia or basidiospore chains were produced by these two double deletion strains (zoomed-in images). All scale bars are 200 μm. (C) Proposed model for the interactions between Ric8 and the three Gα proteins, and how these interactions influence the unisexual development in *C. neoformans*. 24

We hypothesize that the induction of unisexual development in the *ric8*Δ background could be due to mis-regulation of the Gα proteins in the absence of Ric8. To test this hypothesis, we generated all possible single, double, and triple deletion strains of the three Gα proteins in both *MAT*α and *MAT***a** wild-type backgrounds with functional Ric8 and investigated whether they were able to undergo unisexual development under mating inducing conditions (Figure 4B). Of all of the strains tested, only the two independent *MAT*α *gpa2*Δ *gpa3*Δ double deletion strains were able to produce robust hyphae, although they failed to produce basidiospores even after prolonged incubation (>5 weeks). This suggests that both Gpa2 and Gpa3 suppress unisexual development. Interestingly, the two *MAT***a** *gpa2*Δ *gpa3*Δ double deletion strains only grew as yeast cells under mating inducing conditions, similar to all of the other deletion strains tested, suggesting the effects of Gpa2 and Gpa3 on unisexual reproduction could be mating-type dependent. Notably, while the *MAT*α *gpa2*Δ *gpa3*Δ mutant strains produced hyphae, the *MAT*α *gpa1*Δ *gpa2*Δ *gpa3*Δ mutant strains did not, demonstrating that the Gα protein Gpa1 is required for unisexual development in this genetic context even though it was not required on its own in the SSK910 background (deletion of *GPA1* only led to slightly delayed unisexual development in SSK910, Figure 2). We analyzed additional *gpa2*Δ *gpa3*Δ mutant strains constructed in a previous study (53), and found that consistent with the strains generated in this paper, all three *MAT*α strains produced hyphae, while the two *MAT***a** strains did not (Supplemental Figure S7). It should be noted that these deletion strains were constructed in the H99/KN99 background, and as shown earlier, this genetic background is not readily inducible for unisexual developments even when *RIC8* was deleted. Additionally, sexual developments could be influenced by factors such as temperature, moisture, nutrients, and cell density. Thus, it is possible that some of the deletion strains tested could still undergo infrequent unisexual development. Nevertheless, our results suggest that at least some of the Gα proteins function to repress unisexual development under mating inducing conditions, and that modulation of these Gα proteins, likely involving Ric8, could underlie the switch between different modes of sexual development (Figure 4C).

## Discussion

We have discovered that unisexual reproduction can occur in *C. neoformans*, the predominant global pathogenic form of this human fungal pathogen. Our findings further demonstrate that the human fungal pathogen *Cryptococcus* species complex exhibits great plasticity in the ability to achieve successful sexual reproduction, including α-**a** sexual reproduction, unisexual reproduction, and pseudosexual reproduction (4, 11–13, 20). All three modes of sexual reproduction involve hyphal growth, meiosis, and production of basidiospores, which allow the organism to “forage” for nutrients or mating partners over long distances, generate recombinant progeny with potentially higher fitness, and produce basidiospores that are more resilient under challenging conditions and better equipped for dispersal.

We showed that there are variants in the natural population of *C. neoformans* that promote unisexual reproduction. Specifically, while the initial *ric8*11 mutation in the lab strain KN99**a** and KN99α backgrounds only led to sporadic unisexual reproduction, strains SSK110 and SSK910, which are progeny from intercrosses with natural isolates, exhibited significantly enhanced selfing ability. This suggests that it is possible that there are natural strains with enhanced ability to undergo unisexual reproduction. The natural population of *Cryptococcus* species exhibits significant bias towards the α mating type, with populations of more balanced mating types identified only in certain geographic areas (e.g. Sub-Saharan Africa) or certain lineages (e.g. VNBI and VNBII) (17). Thus, the enhanced ability to undergo unisexual reproduction could be favored by natural selection as it allows *Cryptococcus* to undergo sexual development that is beneficial under challenging conditions even in the absence of cells of the opposite mating types required for α-**a** sexual reproduction, and might have emerged as *C. neoformans* became globalized and went through genetic bottlenecks resulting in populations with severely imbalanced mating types. Interestingly, the most self-fertile strain characterized in our study, SSK910, is of the **a** mating type, although the *MAT*α strain SSK110 also exhibits significantly enhanced unisexual fertility. This is consistent with *C. deneoformans*, where unisexual reproduction occurs in both *MAT***a** and *MAT*α strains (54, 55). In *C. deneoformans* the *MAT*α allele has been shown to be a QTL associated with enhances self-filamentation (56), and whether that is also true in *C. neoformans* requires further investigation.

Our findings also highlight the importance of studying strains with diverse genetic backgrounds, including natural strains, to acquire a more complete understanding of biological processes. While studying lab reference strains has its advantages, these isolates could have accumulated private mutations that make them unique compared to other isolates in the species. For example, a recent study showed that the *C. neoformans* reference strain, H99, possesses a unique mutation in the gene *PDE2* that might be contributing to its elevated virulence in the host (57). Another recent study showed that in *C. neoformans* dysfunction in the RNAi pathway leads to hypermutation when combined with and elevated transposon burden in the genome, which was also discovered in natural isolates as the lab strain H99 has a genome with a very low transposon burden (50). Furthermore, the human fungal pathogen *Candida albicans* had been long thought to be RNAi deficient based on the analyses of the reference strains, SC5314, as it contains an inactivating missense mutation in the gene encoding Argonaute, a central RNAi component. However, a recent study of natural *C. albicans* strains showed that the mutation in Argonaute is rare, and the species contains an intact and functional RNAi pathway (58). Thus, what we learn from reference strains may not be representative for the species as a whole.

Our study provides further evidence that *RIC8* is a key regulator of morphological development and a suppressor of unisexual reproduction in *Cryptococcus* species. This is in accord with our previous study in *C. deneoformans* where the presence of a functional *RIC8* restricted the strains to grow as yeast cells under mating inducing conditions, whereas deletion of the *RIC8* gene enabled cells to undergo morphological changes, including hyphal growth and unsexual reproduction, as well as an elevated ability of titan cell formation (27). In *C. deneoformans*, where unisexual reproduction was first characterized, the strain that undergoes the most robust unisexual reproduction, XL280, also contains a nonsense mutation in the *RIC8* gene (27). Additionally, during our effort to construct *C. neoformans* strains that undergo robust unisexual reproduction, all of the selfing progeny inherited the *ric8*11 deletion allele. Thus, *RIC8* appears to be a critical regulator of morphological development in *Cryptococcus* species. The *RIC8* gene encodes a guanine nucleotide exchange factor (GEF) for the Gα subunit of heterotrimeric G proteins, and our analyses indicate that it structurally may form protein complexes with all three Gα proteins similarly. Ric8 could function through modulating G-protein coupled receptor (GPCR) signaling pathways, by facilitating GDP-to-GTP exchange and activating Gα subunits, as well as serving as a chaperone for the Gα proteins, which in turn interact with downstream effectors and initiate various intracellular signaling pathways (59, 60). It is possible that under mating inducing conditions, Ric8 activates Gα proteins (Gpa1, Gpa2, and maybe Gpa3 as well) that favors α-**a** sexual reproduction and represses unisexual reproduction, which is also consistent with the observation that *ric8*11 strains exhibit significantly reduced mating in α-**a** sexual reproduction (27), providing selective pressure for the conservation of the *RIC8* gene. In contrast, the absence of Ric8 could lead to changes in the modulation of the Gα proteins (e.g. activity and localization), enabling cells to initiate sexual reproduction without the presence of compatible mating types when conditions become favorable for mating, which could be through modulation of cell ploidy (haploid to diploid) and/or morphology (yeast to hyphae). Previous studies have also identified and characterized various other roles of the three Gα subunits in *C. neoformans* (Gpa1, Gpa2, and Gpa3).

Specifically, Gpa1 physically and functionally interacts with the nutrient-sensing GPCR Gpr4 and functions to activate the cAMP-PKA signaling pathway that controls sexual reproduction, virulence factor production, and pathogenesis (45, 61); Gpa2 and Gpa3 are both required for pheromone sensing and sexual reproduction (e.g. *gpa2*11 *gpa3*11 double mutants are sterile but *gpa2*11 or *gpa3*11 mutants are not), whereas loss of Gpa3 leads to constitutive activation of the pheromone response pathway in *C. deneoformans*, illustrating an inhibitory role (53). Because Gα proteins can play roles in governing signal transduction in both their GDP inactive and GTP active state, further studies will be required to elucidate the precise organization of the Ric8 controlled Gα protein signaling networks, as well as how the switching between α-**a** sexual reproduction and unisexual reproduction is regulated by *RIC8* and the Gα proteins, and whether this regulatory pathway is conserved in other fungal species.

Different modes of sexual reproduction in *Cryptococcus* involve shared molecular pathways, including a functional MAPK pathway for signal transduction; transcription factors that play central roles in regulating morphological development during sexual reproduction (e.g. Mat2 and Znf2); and key factors such as Dmc1 and Spo11 for meiosis and sporulation. Thus, the difference mainly lies in how sexual reproduction is initiated. It is known that certain environmental cues, including certain nutrients and temperature, induce sexual reproduction in *Cryptococcus*. Additionally, sexual reproduction in *Cryptococcus* may also be a response to biotic factors in the environment, such as fungal/bacterial cohabitants and their secondary metabolites, which can induce stress response, lead to transcriptome fluctuation, and modulate morphological changes, and for which many of the response pathways, including MAPK, cAMP-PKA, calcineurin, and TOR pathways, have been shown to be involved in regulating morphogenesis and sexual development in *Cryptococcus* (62).

As a unicellular eukaryote, *Cryptococcus* is sensitive to environmental changes. Being able to initiate diverse modes of sexual reproduction, α-**a** sexual reproduction when both mating types are present and unisexual reproduction when only one mating type is available, allows *Cryptococcus* to grow by mitotic propagation under favorable environmental conditions, and rapidly undergo sexual reproduction when challenging environmental conditions arise. It is possible that the Ric8-mediated regulatory pathway for unisexual reproduction is conserved and modulated by yet to be characterized environmental factors and conditions, through modulation of the Ric8 activity and stability. Unisexual reproduction may occur in other fungi, shaping the ecology, population dynamics, and evolutionary trajectory of not only the pathogenic *Cryptococcus* species, but also other fungal species and possibly eukaryotic microorganisms beyond the fungal kingdom.

## Materials and methods

### Strain handling and culturing

All of the strains were grown on YPD medium, supplemented with appropriate antifungal drugs (nourseothricin (NAT) at 100 μg/mL or neomycin (G418) at 200 μg/mL) when needed, and incubated at 30 °C unless stated otherwise.

### DNA preparation and whole genome sequencing

DNA for genotyping and Illumina whole genome sequencing were prepared using a MasterPure Yeast DNA Purification Kit (LGC Biosearch Technologies, MPY80200) as previously described (63). Illumina sequencing was conducted at the Duke Sequencing and Genomic Technologies core facility (https://genome.duke.edu), on a Novaseq 6000 platform with 250 bp paired-end option.

High molecular weight genomic DNA samples for Nanopore long read sequencing were prepared using a modified CTAB protocol as previously described (4). Nanopore long-read sequencing was conducted in house using a MinION device with R10.4.1 flow cells following instructions provided by the manufacturer.

All of the sequencing data have been deposited in the BioProject ID PRJNA1269048.

### Deletion strain construction

Gene deletions using the *NAT* or *NEO* dominant markers were conducted using a TRACE CRISPR system (64) or biolistic transformation (*GPA3*, *STE3***a**, and *STE11***a**). For CRISPR deletion of each gene of interest, one or two guide RNA was used to introduce double strand breaks within the gene, and the open reading frame (ORF) was then replaced through homologous recombination using either a *NAT* or a *NEO* deletion allele that contains a drug-resistant gene cassette at the center, flanked on both side by sequences (∼1 kb) homologous to the 5’- and 3’-regions adjacent to the target gene ORF. For biolistic transformation, similar deletion allele was used following the protocol published previously (65). The obtained transformants were validated by genotyping using diagnostic PCRs for the internal region of the ORF, 5’-junction and 3’-junction regions, as well as spanning PCR of the region encompassing the deletion construct as previously described (66) (Supplemental Figure S5). The information on the guide RNAs, as well as primers used to build deletion constructs and validate transformants, are included in the Supplemental Table S3.

### Mating and spore dissection

The mating assay and spore dissection were carried as described in previously published protocols (67). Briefly, all of the mating assays were conducted on MS solid medium at room temperature in the dark. Plates were inspected for sexual development after 1-2 weeks of incubation and kept for prolonged incubation when needed. Basidiospore dissection was carried out using a Nikon yeast spore dissection microscope equipped with a glass fiber needle 25 μm in diameter.

### Flow cytometry analysis

Samples for flow cytometry analysis to determine *Cryptococcus* ploidy were prepared as previously described (63, 68), and then submitted to and analyzed by the Duke Cancer Institute Flow Cytometry Shared Resource Laboratory.

### Microscopy imaging

Brightfield and differential interference contrast (DIC) microscopy images were acquired with an AxioScop 2 fluorescence microscope equipped with an AxioCam MRm digital camera (Zeiss, Germany). Scanning electron microscopy (SEM) was performed at the Duke Microscopy Core Facility using a scanning electron microscope with an EDS detector (Apreo S, ThermoFisher, USA).

## Acknowledgements

We thank Dr. Steve Haase for providing constructive feedback that helped improve the manuscript. This work was supported by NIH/NIAID awards R01AI039115-28, R01AI050113-20, R01AI170543-04, R01AI172451-03, and R01AI133654-08 to JH. JH is a fellow and co-director with Leah Cowen of the CIFAR program Fungal Kingdom: Threats & Opportunities.

## Statement of competing interest

The authors declare no competing interests.

**Supplemental Figure S1. Genotypic profiles of the unisexually fertile *C. neoformans* strains**. (A) Genotyping of the *MAT* locus for the haploid strains analyzed in this study. (B) Genome analysis of the strains that unisexually reproduce. The blue and red colors indicate SNPs when mapped against the H99 and Bt63 genomes, respectively. For each strain, zoomed-in read coverage maps of chromosome 5 are shown with. the H99 (blue) or Bt63 (red) genome as reference. For strains SSK110 and SSK910, additional zoomed-in maps of SNPs against H99 (blue) or Bt63 (red) along chromosome 5 are included beneath the read-depth plots.

**Supplemental Figure S2. Analyses of progeny from diploid *MAT*a/a *C. neoformans* strain SSL570.**

Variant mapping of the parental strain SSL570 and its dissected basidiospore chains. For the parental strain SSL570, the variants against H99 (blue) and Bt63 (red) were mapped against the H99 and Bt63 genomes, respectively. For the progeny, the variants against H99 and Bt63 were all mapped with the H99 genome as reference. The purple column indicates the location of the *MAT* locus.

**Supplemental Figure S3. Analyses of progeny from diploid *MAT*α/α *C. neoformans* strain SSL571.**

Variant mapping of the parental strain SSL571 and its dissected basidiospore chains. For the parental strain SSL571, the variants against H99 (blue) and Bt63 (red) were mapped against the H99 and Bt63 genomes, respectively. For the progeny, the variants against H99 and Bt63 were all mapped with the H99 genome as reference. The purple column indicates the location of the *MAT* locus.

**Supplemental Figure S4. Analyses of progeny from diploid *MAT*α/α *C. neoformans* strain SSL576.**

Variant mapping of the parental strain SSL576 and its dissected basidiospore chains. For the parental strain SSL576, the variants against H99 (blue) and Bt63 (red) were mapped against the H99 and Bt63 genomes, respectively. For the progeny, the variants against H99 and Bt63 were all mapped with the H99 genome as reference. The purple column indicates the location of the *MAT* locus.

**Supplemental Figure S5. PCR validation of the deletion strains in the self-fertile SSK910 strain background.**

Each image demonstrates PCR validation of two independent deletion strains of one gene (labeled at the right bottom corner of the image), including 5’- and 3’-junction PCRs (left and right panels, respectively, amplifying only from deletion strains) and internal PCR (middle panel, amplifying only from wild-type controls). For each PCR reaction, samples from left to right are: deletion strain No.1, deletion strain No.2, KN99a wild-type control, KN99α wild-type control, and ddH2O negative PCR control.

**Supplemental Figure S6. Selfing assay of mutant strains in the self-fertile *C. neoformans* SSK910 strain background.**

Light microscopy images of strains with deletions of various genes in the SSK910 background when induced for unisexual reproduction on MS medium. Images shown are from two independent deletion strains of each gene, with images on the right showing the basidia with or without typical basidiospore chains for the deletion strains that were still able to undergo unisexual reproduction. Scales are shown in the images of the SSK910 progenitor strain.

**Supplemental Figure S7. Selfing assay of gpa2Δ gpa3Δ mutant strains.**

(A) The five *gpa2*11 *gpa3***Δ** mutant strains constructed in a previous study (53). (B) Selfing assay of the five *gpa2***Δ** *gpa3***Δ** mutant strains under mating inducing condition. Images were taken after 10 days of incubation.

---

**Supplemental Table S1.** Strains analyzed in this study.

| Strain Name | Strain background | Genotype | Source |
| --- | --- | --- | --- |
| KN99α | n.a. | *MAT*α wildtype | (70) |
| KN99**a** | n.a. | *MAT***a** wildtype | (70) |
| AD1-7a | n.a. | *MAT*α wildtype | (19) |
| T4 | n.a. | *MAT*α wildtype | (19) |
| Bt63 | n.a. | *MAT***a** wildtype | (19) |
| Bt65 | n.a. | *MAT***a** wildtype | (19) |
| SSI867 | KN99α | *MAT*α *ric8*Δ::*NAT* | (27) |
| SSI868 | KN99α | *MAT*α *ric8*Δ::*NAT* | (27) |
| SSI869 | Progeny from SSI867 × KN99**a** | *MAT***a** *ric8*Δ::*NAT* | (27) |
| SSI870 | Progeny from SSI868 × KN99**a** | *MAT***a** *ric8*Δ::*NAT* | (27) |
| SSK110 | Progeny from SSI869 × AD1-7a | *MAT*α *ric8*Δ::*NAT* recombinant genome | This study |
| SSK910 | Progeny from SSK110 × Bt63 | *MAT***a** *ric8*Δ::*NAT* recombinant genome | This study |
| SSL436 | Bt63 | *MAT***a** *ric8*Δ::*NEO* | This study |
| SSL570 | SSL436 + SSK110 fusion No.06 | *MAT***a/a** *NAT NEO* | This study |
| SSL571 | SSL436 + SSK110 fusion No.07 | *MAT*$\alpha/\alpha$ *NAT NEO* | This study |
| SSL576 | SSL436 + SSK110 fusion No.12 | *MAT*$\alpha/\alpha$ *NAT NEO* | This study |
| SSL739 | SSK910 | *MAT***a** *gpa2*Δ::*NEO*_1 | This study |
| SSL740 | SSK910 | *MAT***a** *gpa2*Δ::*NEO*_2 | This study |
| SSL741 | SSK910 | *MAT***a** *pka1*Δ::*NEO*_1 | This study |
| SSL742 | SSK910 | *MATa pka1Δ::NEO_2* | This study |
| SSL743 | SSK910 | *MATa ste7Δ::NEO_1* | This study |
| SSL744 | SSK910 | *MATa ste7Δ::NEO_2* | This study |
| SSL745 | SSK910 | *MATa cpk1Δ::NEO_1* | This study |
| SSL746 | SSK910 | *MATa cpk1Δ::NEO_2* | This study |
| SSL747 | SSK910 | *MATa znf2Δ::NEO_1* | This study |
| SSL748 | SSK910 | *MATa znf2Δ::NEO_2* | This study |
| SSL749 | SSK910 | *MATa cpa1Δ::NEO_1* | This study |
| SSL750 | SSK910 | *MATa cpa1Δ::NEO_2* | This study |
| SSL751 | SSK910 | *MATa gpr4Δ::NEO_1* | This study |
| SSL752 | SSK910 | *MATa gpr4Δ::NEO_2* | This study |
| SSL753 | SSK910 | *MATa mat2Δ::NEO_1* | This study |
| SSL754 | SSK910 | *MATa mat2Δ::NEO_2* | This study |
| SSL755 | SSK910 | *MATa sxi2Δ::NEO_1* | This study |
| SSL756 | SSK910 | *MATa sxi2Δ::NEO_2* | This study |
| SSL757 | SSK910 | *MATa spo11Δ::NEO_1* | This study |
| SSL758 | SSK910 | *MATa spo11Δ::NEO_2* | This study |
| SSL759 | SSK910 | *MATa dmc1Δ::NEO_1* | This study |
| SSL760 | SSK910 | *MATa dmc1Δ::NEO_2* | This study |
| YSB25 | H99 | *MATα gpa2Δ::NEO* | (53) |
| YSB26 | H99 | *MATα gpa2Δ::NEO* | (53) |
| YSB85 | KN99a | *MATa gpa1Δ::NEO* | (71) |
| YSB86 | KN99a | *MATa gpa1Δ::NEO* | (71) |
| YSB137 | KN99a | *MATa gpa3Δ::NEO* | (53) |
| YSB138 | KN99a | *MATa gpa3Δ::NEO* | (53) |
| SSL808 | YSB25 × YSB85 | *MATα gpa1Δ::NEO* | This study |
| SSL809 | YSB25 × YSB85 | *MATα gpa1Δ::NEO* | This study |
| SSL810 | YSB25 × YSB85 | *MATa gpa2Δ::NEO* | This study |
| SSL811 | YSB25 × YSB85 | *MATa gpa2Δ::NEO* | This study |
| SSL813 | YSB25 × YSB85 | *MATα gpa1Δ::NEO gpa2Δ::NEO* | This study |
| SSL832 | SSL813 × YSB137 | *MATα gpa3Δ::NEO* | This study |
| SSL833 | SSL813 × YSB137 | *MATα gpa3Δ::NEO* | This study |
| SSL815 | SSL813 × YSB137 | *MATa gpa1Δ::NEO gpa2Δ::NEO* | This study |
| SSL816 | SSL813 × YSB137 | *MATa gpa1Δ::NEO gpa2Δ::NEO* | This study |
| SSL817 | SSL813 × YSB137 | *MATα gpa1Δ::NEO gpa2Δ::NEO* | This study |
| SSL818 | SSL813 × YSB137 | *MATα gpa1Δ::NEO gpa2Δ::NEO* | This study |
| SSL819 | SSL813 × YSB137 | *MATa gpa1Δ::NEO gpa3Δ::NEO* | This study |
| SSL820 | SSL813 × YSB137 | *MATa gpa1Δ::NEO gpa3Δ::NEO* | This study |
| SSL821 | SSL813 × YSB137 | *MATα gpa1Δ::NEO gpa3Δ::NEO* | This study |
| SSL822 | SSL813 × YSB137 | *MATα gpa1Δ::NEO gpa3Δ::NEO* | This study |
| SSL823 | SSL813 × YSB137 | *MATa gpa2Δ::NEO gpa3Δ::NEO* | This study |
| SSL824 | SSL813 × YSB137 | *MATa gpa2Δ::NEO gpa3Δ::NEO* | This study |
| SSL825 | SSL813 × YSB137 | *MATα gpa2Δ::NEO gpa3Δ::NEO* | This study |
| SSL831 | SSL813 × YSB137 | *MATα gpa2Δ::NEO gpa3Δ::NEO* | This study |
| SSL827 | SSL813 × YSB137 | *MATa gpa1Δ::NEO gpa2Δ::NEO gpa3Δ::NEO* | This study |
| SSL828 | SSL813 × YSB137 | *MATa gpa1Δ::NEO gpa2Δ::NEO gpa3Δ::NEO* | This study |
| SSL829 | SSL813 × YSB137 | *MATα gpa1Δ::NEO gpa2Δ::NEO gpa3Δ::NEO* | This study |
| SSL830 | SSL813 × YSB137 | *MATα gpa1Δ::NEO gpa2Δ::NEO gpa3Δ::NEO* | This study |
| YPH106 |  | *MATa gpa2Δ::NEO gpa3Δ::NEO* | (53) |
| YPH118 |  | *MATa crg1Δ::URA5 gpa2Δ::NEO gpa3Δ::NEO* | (53) |
| YPH305 |  | *MATα crg2Δ::NAT gpa2Δ::NEO gpa3Δ::NEO* | (53) |
| YPH308 |  | *MATα gpa2Δ::NAT gpa3Δ::NEO* | (53) |
| YPH380 |  | *MATα crg1Δ::URA5 gpa2Δ::NEO gpa3Δ::NEO* | (53) |

**Supplemental Table S2.** Genes deleted in the self-fertile strain.

| Gene ID <sup>1</sup> | Gene Name | Encoded protein | Self-fertility | Sporulation |
| --- | --- | --- | --- | --- |
| CNAG_04505 | *GPA1* | Guanine nucleotide-binding protein subunit $\alpha$ | Slightly delayed | Yes |
| CNAG_00179 | *GPA2* | G protein $\alpha$ subunit | Yes | Yes |
| CNAG_02090 | *GPA3* | G protein $\alpha$ subunit | Yes | Yes |
| CNAG_06808 <sup>2</sup> | *STE3* | G-protein coupled receptor | Yes | Yes |
| CNAG_04730 | *GPR4* | G-protein coupled receptor | Yes | Yes |
| n.a. <sup>1</sup> | *SXI2* | Homeodomain (HD) transcription factor | Yes | Yes |
| CNAG_00396 | *PKA1* | Protein kinase A | Slightly delayed | Yes |
| CNAG_05472 | *SPO11* | Meiotic recombination protein | Yes | No |
| CNAG_07909 | *DMC1* | Meiotic recombinase | Yes | No |
| CNAG_02511 | *CPK1* | MAPK | No | n.a. |
| CNAG_01730 | *STE7* | MAPKK | No | n.a. |
| CNAG_06980 <sup>2</sup> | *STE11* | MAPKKK | No | n.a. |
| CNAG_06203 | *MAT2* | HMG-box transcription factor | No | n.a. |
| CNAG_03366 | *ZNF2* | Master regulator of filamentation, C2H2 type zinc finger | No | n.a. |

<sup>1</sup>: The gene IDs are based on the annotation of the *MAT $\alpha$* H99 genome. *SXI2a* does not have a gene ID as it is a *MATa* specific gene.
<sup>2</sup>: *STE3* and *STE11* are mating-type specific genes, and their *MATa* homologs were deleted in the *MATa* self-fertile strain SSK910.

Supplemental Table S3. Primers used in this study.

(Please see the attached Excel file)

