## Supplementary figures and images for "Unisexual reproduction in the global human fungal pathogen *Cryptococcus neoformans*"

### Supplemental Figure S1

Supplemental Figure S1

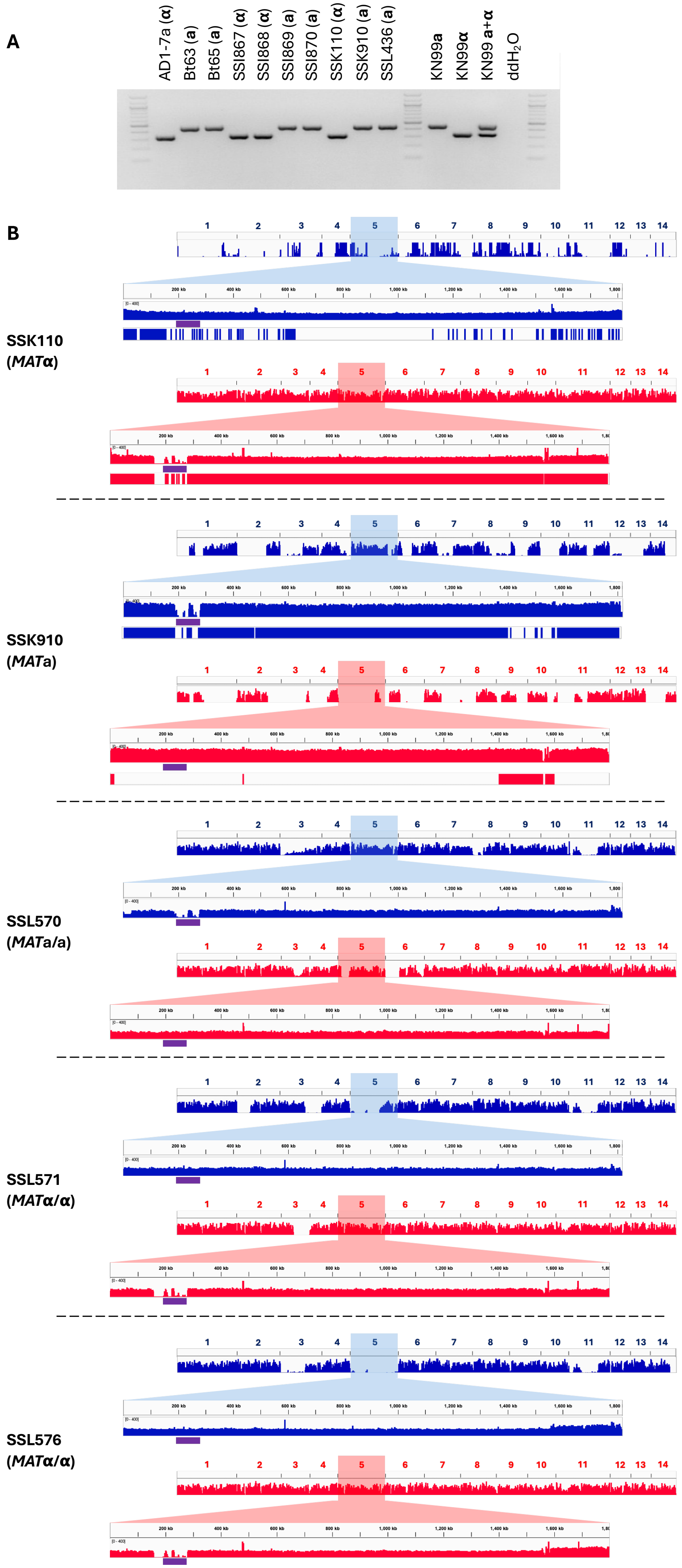

### Supplemental Figure S2

#### Supplemental Figure S2

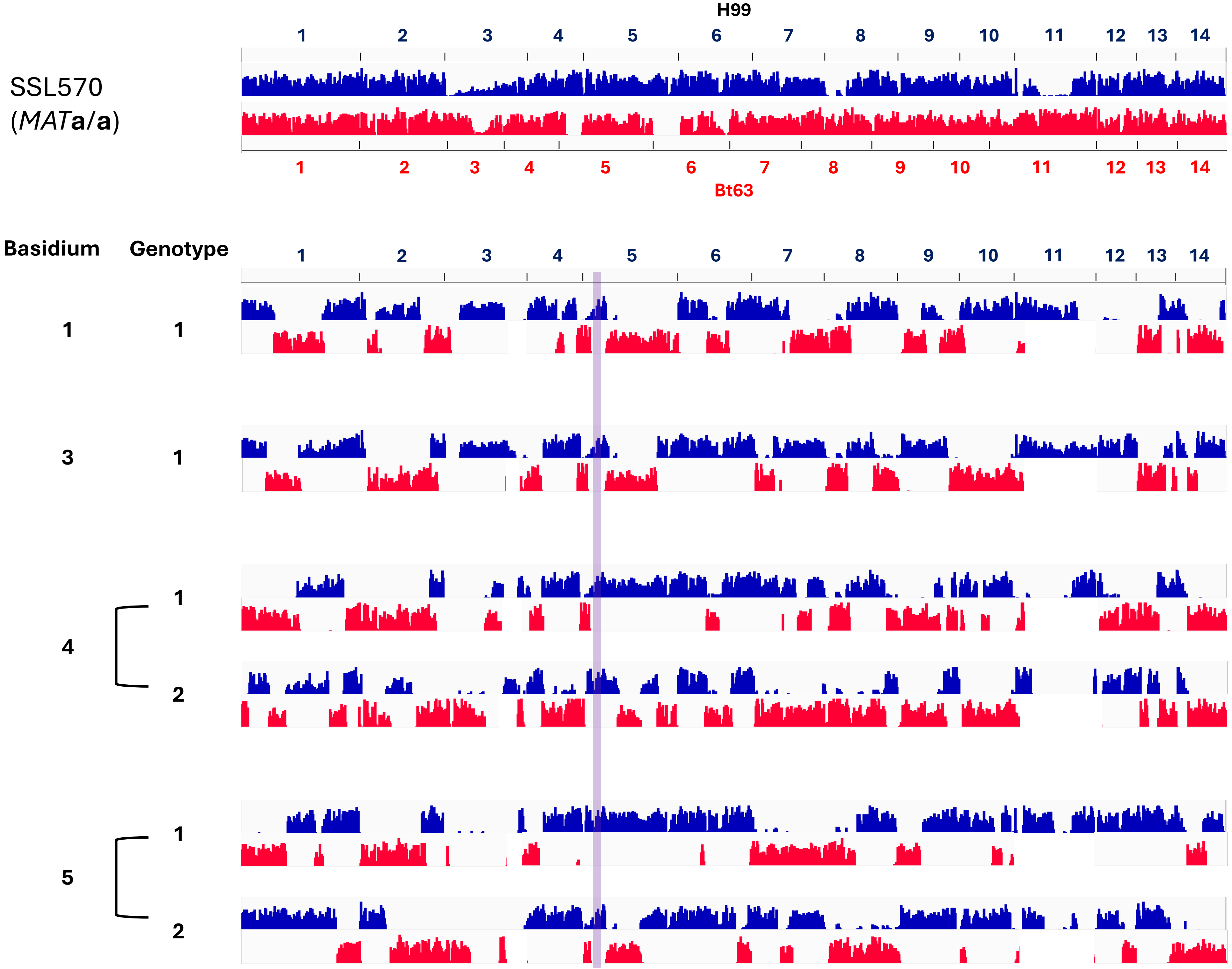

### Supplemental Figure S3

#### Supplemental Figure S3

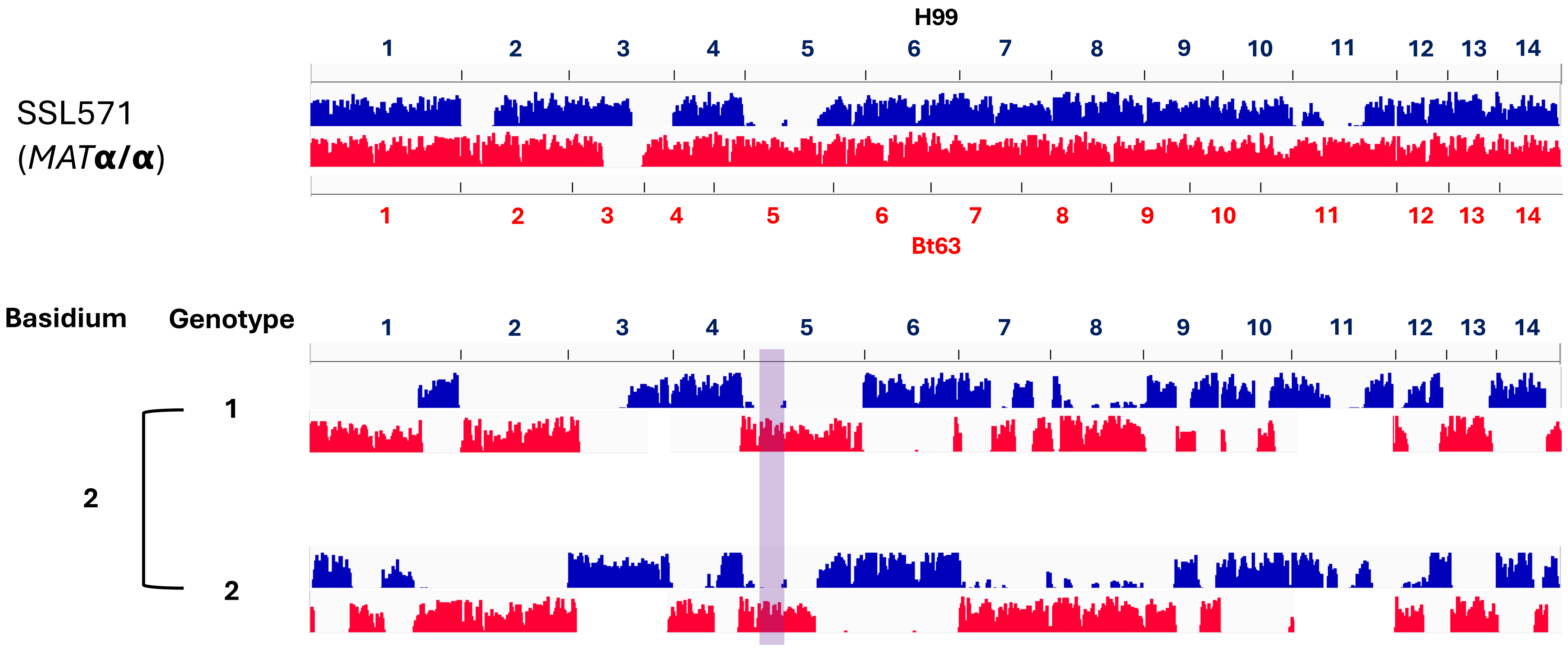

### Supplemental Figure S4

**Supplemental Figure S4**

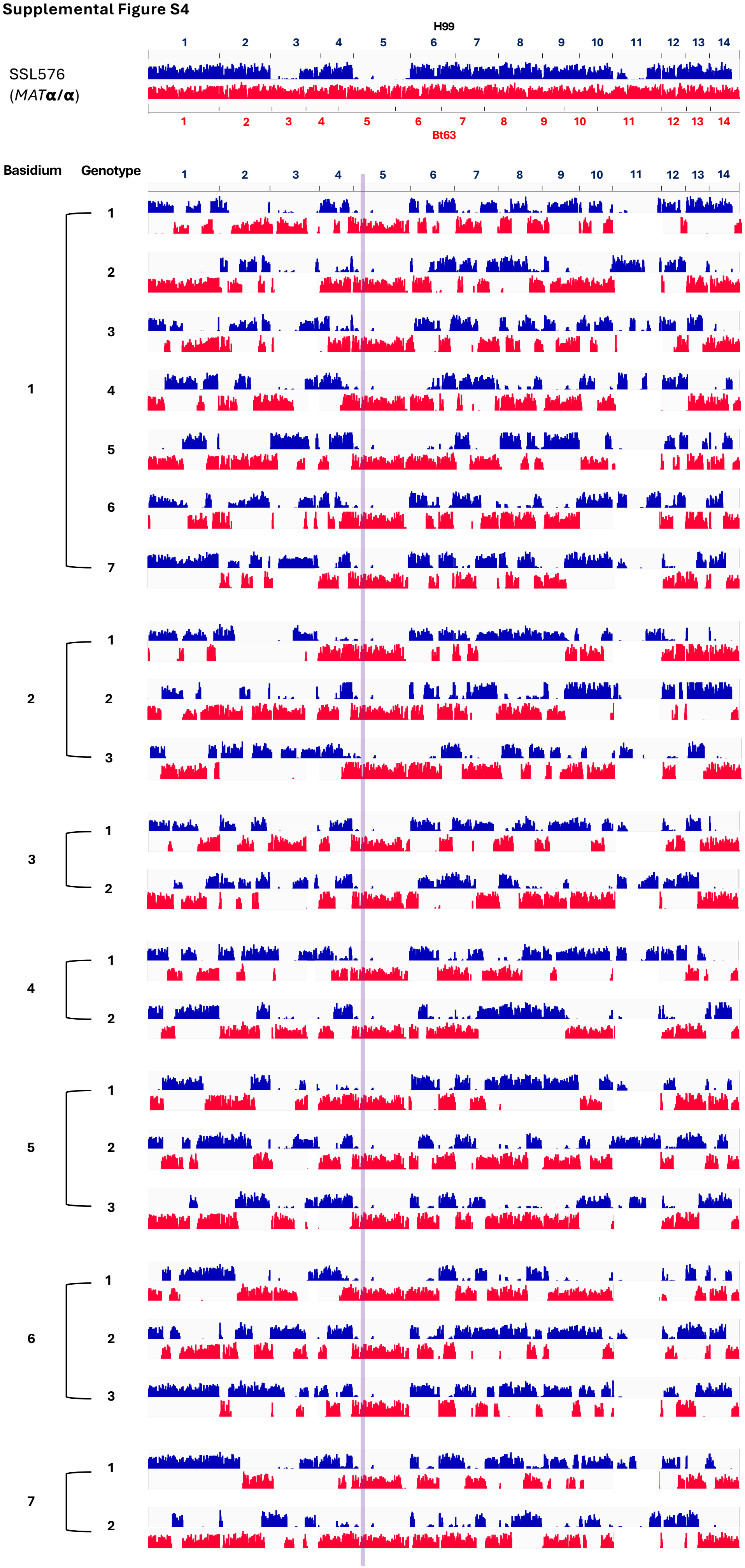

### Supplemental Figure S6

#### Supplemental Figure S6

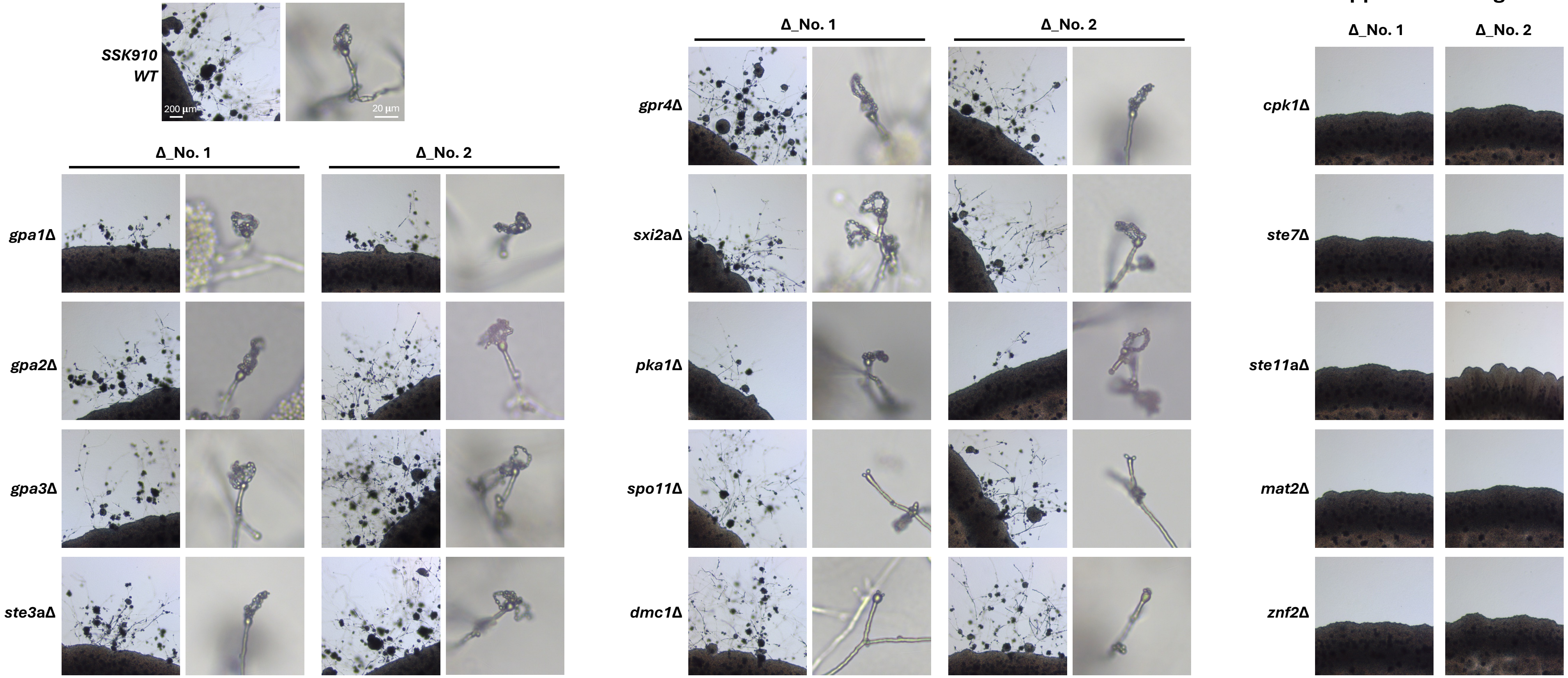
