## Supplemental Figure S5 for "Unisexual reproduction in the global human fungal pathogen *Cryptococcus neoformans*"

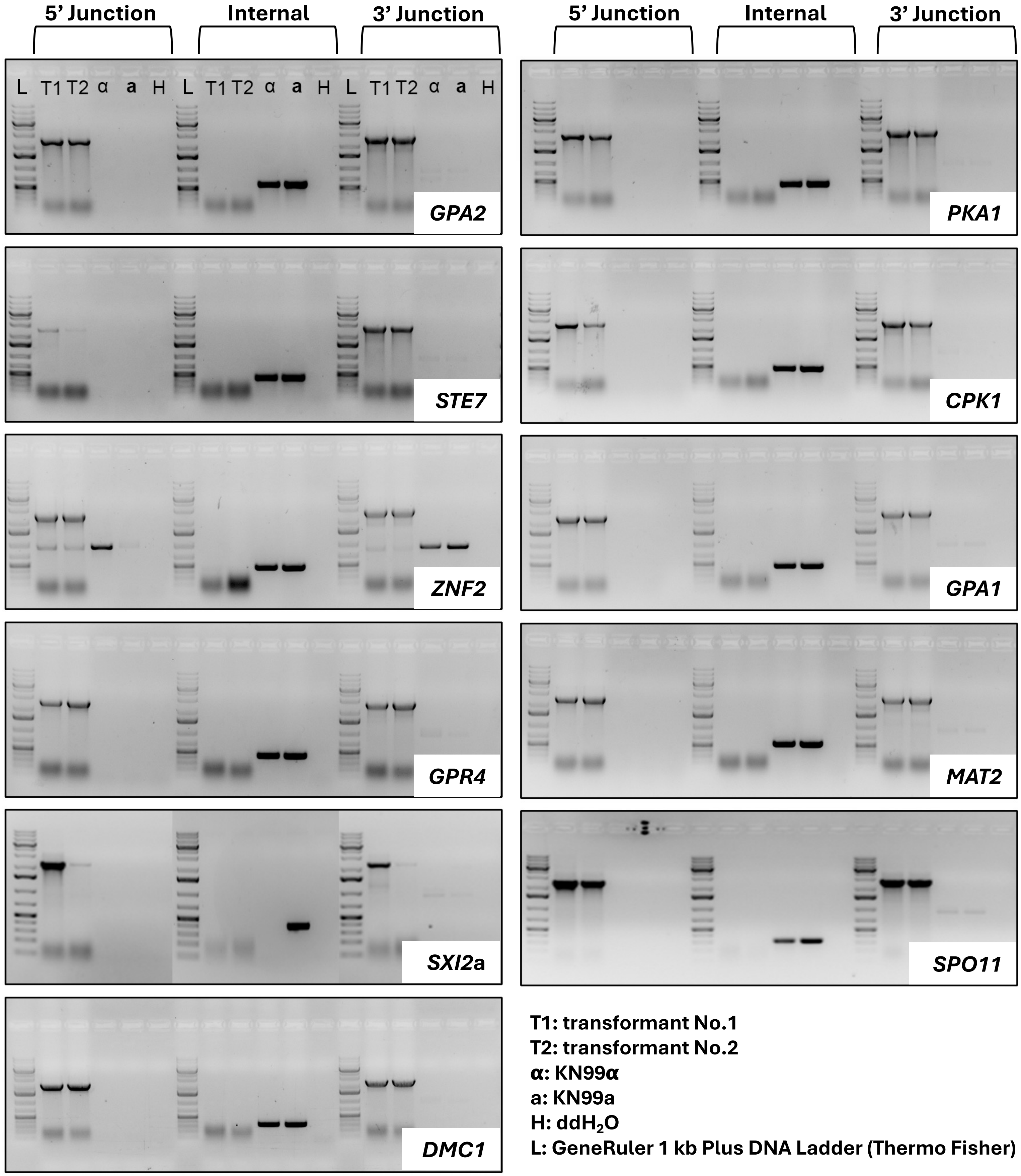


**T1: transformant No.1**
**T2: transformant No.2**
**α: KN99α**
**a: KN99a**
**H: dd$H_2O$**
**L: GeneRuler 1 kb Plus DNA Ladder (Thermo Fisher)**
