## Supplemental Figure S7 for "Unisexual reproduction in the global human fungal pathogen *Cryptococcus neoformans*"

**A**

| Strain Name | Mating Type | Genotype / Markers / Phenotype |
|---|---|---|
| YPH106 | **a** | *gpa2Δ::NEO gpa3Δ::NEO* |
| YPH118 | **a** | *crg1Δ::URA5 gpa2Δ::NEO gpa3Δ::NEO* |
| YPH305 | *α* | *crg2Δ::NAT gpa2Δ::NEO gpa3Δ::NEO* |
| YPH308 | *α* | *gpa2Δ::NAT gpa3Δ::NEO* |
| YPH380 | *α* | *crg1Δ::URA5 gpa2Δ::NEO gpa3Δ::NEO* |

**B**

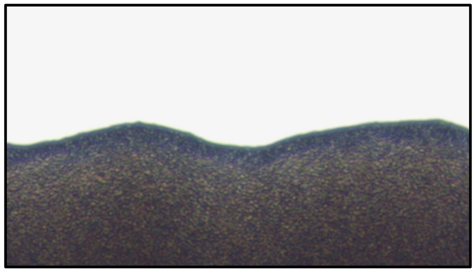

**YPH106**

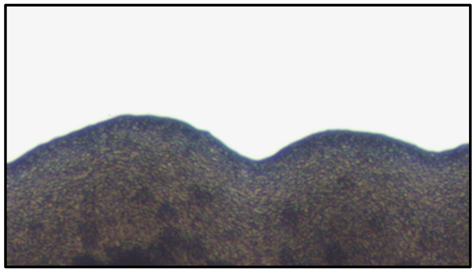

**YPH118**

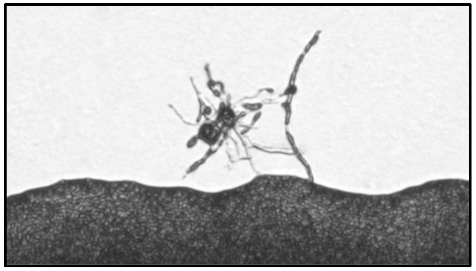

**YPH305**

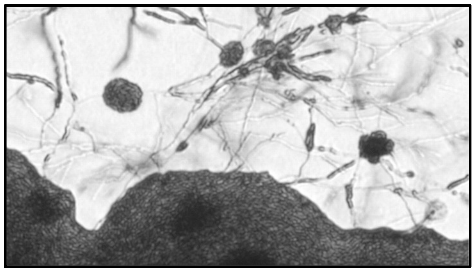

**YPH308**

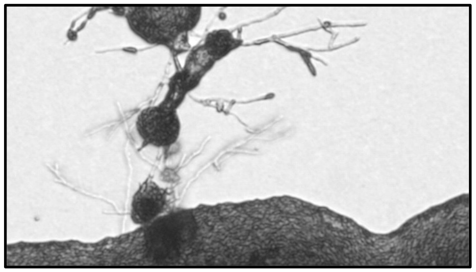

**YPH380**
